# Systems modeling identifies phenotype-determining signaling pathways controlled by phosphatase PTPRJ in diverse receptor tyrosine kinase activation settings

**DOI:** 10.64898/2026.04.30.721884

**Authors:** Will S. Hart, Katie M. Knight, Sophie Rizzo, Sung Hyun Lee, Rachel Fetter, Damien Thevenin, Matthew J. Lazzara

## Abstract

Protein tyrosine phosphatase receptor J (PTPRJ) restrains cell proliferation and migration by dephosphorylating receptor tyrosine kinases (RTKs) including the epidermal growth factor receptor (EGFR). PTPRJ is a purported tumor suppressor, and alterations to its expression and/or function are associated with colorectal, breast, lung, and other cancers. While there is interest in controlling PTPRJ-regulated phenotypes, efforts are limited by the complexity of PTPRJ-mediated signaling. PTPRJ dephosphorylates multiple RTKs, and the degree to which PTPRJ control of signaling and phenotypes depends on local cellular RTK activation profiles is unknown. To probe the context dependence of PTPRJ signaling regulation, we collected signaling measurements across 16 pathway nodes at two time points in a panel of HSC3 carcinoma cells engineered with different PTPRJ expression profiles. Cells were treated with three different RTK ligands, and paired phenotype measurements (viability, wound healing, xCELLigence cell index) were made. Partial least squares regression models were developed to predict relationships between PTPRJ-regulated signaling pathways and cell phenotypes. The model effectively separated contributions to variance arising from the PTPRJ expression background and growth factor context. In testing model predictions, we demonstrated that PTPRJ suppressed MET-induced cell cell proliferation via regulation of a HER3/AKT signaling axis that stabilized PTPRJ expression through an unanticipated feedback mechanism. We also found that PTPRJ regulated HSC3 cell migration via JNK signaling that was preferentially activated by MET. Our results identify new regulatory nodes through which PTPRJ influences cancer cell phenotypes and demonstrates that these processes preferentially occur in the context of distinct RTK activation states.

## INTRODUCTION

Protein tyrosine phosphatase receptor J (PTPRJ), also known as density-enhanced phosphatase 1 (DEP1), is a receptor-like protein tyrosine phosphatase (RPTP) and purported tumor suppressor. PTPRJ loss of heterozygosity (LOH) occurs in colorectal (1) and breast cancers (2, 3), and PTPRJ polymorphisms have been observed in non-Hodgkin’s lymphoma (4), breast cancer (5), and thyroid cancer (6). The R326Q SNP is associated with increased colorectal cancer risk (3, 7), but the signaling effects of this mutation have not been characterized. The naturally occurring Q276P and R326Q extracellular domain mutations do not alter signaling (8). Knockout or knockdown of PTPRJ controls cancer-relevant phenotypes including migration and proliferation (9, 10), and silenced PTPRJ expression through promoter hypermethylation has been observed in patient-derived tumors (11).

Many potential PTPRJ-regulated substrates have been reported, and substrate-trapping mutants (STMs) of PTPRJ have been engineered (12). These catalytically inactive mutants (D1205A or C1239S) stably complex with potential substrates that can be identified through immunoprecipitation followed by immunoblot or mass spectrometry. PTPRJ STMs bind receptor tyrosine kinases (RTKs) including epidermal growth factor receptor (EGFR) (12), MET (13–15), Fms-related receptor tyrosine kinase (FLT3) (16), insulin receptor (INSR) (17), and platelet-derived growth factor receptor (PDGFR) (18). PTPRJ STMs also bind non-receptors including the p85 regulatory subunit of PI3K (19), ERK1/2 (20), GAB1 (13), and p120-catenin (13, 14). PTPRJ also indirectly regulates proteins that have not been reported to be captured by STMs, including vascular endothelial growth factor receptor (VEGFR) (21, 22), AKT (23, 24), and JAK2 (25). Given the vast array of signaling nodes regulated by PTPRJ, it has been difficult to elucidate which aspects of its function are most important for controlling cell phenotypes.

The catalytic activity of PTPRJ and other RPTPs is regulated by oligomerization. Unlike the regulation of classical RTKs (26), oligomerization inhibits RPTP activity (27–29). We previously showed that PTPRJ dimerization is regulated by key glycine residues in the transmembrane domain (TMD) and that mutating these residues to bulky ones (e.g., G983L) destabilizes oligomerization, promoting an active, monomeric state of PTPRJ (29). Oligomer-destabilizing PTPRJ mutations decrease EGFR and FLT3 phosphorylation and carcinoma-associated phenotypes (28, 29), and peptide agonists targeting the TMD domain inhibit EGFR-driven proliferation and migration (29, 30).

Even with substantial characterization of proteins regulated by RTK-RPTP interactions (31), the complexity of the RPTP-controlled network dynamics and network control of phenotypes is far from fully understood. Computational modeling has been leveraged to address some of the gaps. For example, mechanistic modeling has been used to infer rates of EGFR dephosphorylation at the plasma membrane and endosome (32), explain signaling complex persistence distal from EGFR through repeated phosphorylation and dephosphorylation cycles (33), understand the control of time-varying EGFR signaling by receptor recycling (34), and gain insight into the ability of EGFR ubiquitination to control a switch between suppression of spurious and ligand-induced signaling (35). Mechanistic models are especially useful for elucidating early RPTP- or receptor-proximal aspects signaling regulation, where model identifiability is reasonable, but it is generally challenging for mechanistic models to lend insight into the multivariate signaling control of cell phenotypes (36). Machine learning models based on regression, classification, and other methods have become increasingly common as the rate of biomedical data generation has exploded (37), and the application of these methods has been seen in studies of signaling. For example, partial least squares regression (PLSR) was used to decipher the key HER2-regulated tyrosine phosphorylation events that control mammary epithelial cell fates (38), and integrated mechanistic-machine learning models have been used to predict cell phenotypic responses to different EGFR ligands (39). To our knowledge, only one prior study has applied machine learning to identify the phenotype-controlling signaling pathways regulated by a phosphatase (40). The results of that study clearly demonstrated the ability of machine learning to identify unanticipated regulation of signaling and phenotypes by SHP2.

Here, we used network-level phosphoprotein measurements to quantify the effects of PTPRJ on signaling and phenotype regulation in a human squamous carcinoma cell line treated with three different growth factors. Partial least squares regression (PLSR) data-driven modeling was employed to predict the relationship between the signaling nodes and phenotypes including viability, wound healing, and xCELLigence cell index. The model nominated the HER3 psuedokinase as a key PTPRJ-regulated node that controls cell index and pointed to JNK and CREB as PTPRJ-regulated signaling nodes altering the wound-healing phenotype. HGF perturbed these phenotypes the most, and PTPRJ’s role as a phosphatase in altering phenotypic outcomes was growth factor dependent. We also observed an unexpected change in ectopically expressed PTPRJ protein levels in response to HER3 knockdown and identified a novel PTPRJ-ErbB receptor feedback loop.

## METHODS

### Cell culture

Human oral squamous carcinoma (HSC3) cells (Alexander Sorkin, University of Pittsburgh), which have low endogenous PTPRJ expression, were cultured in high-glucose DMEM (Gibco 11965118) supplemented with 10% fetal bovine serum (Avantor 97068-085; lot number 256K22), 100 units/mL penicillin (ThermoFisher 15140122), 0.1 mg/mL streptomycin (ThermoFisher 15140122), and 1 mM L-glutamine (ThermoFisher 25030081) in a humidified ThermoForma i160 incubator at 5% CO_2_ and 37°C.

### Cell line engineering

HSC3 cells were transduced with pBABE-puro retroviral vectors encoding wild-type (WT) or oligomerization-impaired G983L mutant PTPRJ, or with the empty vector (EV) as a control. The creation of these stable transductants was previously described (29). Transduced populations and clonal lines derived from them were maintained in 1 µg/mL puromycin (GeminiBio 400-128P-100).

### Generation of clonal lines of HSC3 transductants

EV, WT, G983L HSC3 transductants were single-cell sorted at the UVA Flow Cytometry Core Facility in 96-well plates. Clones were expanded, frozen, and screened by western blotting to identify WT and G983L clones with similar EGFR and PTPRJ expression. The clones used were: P4D4 (EV), P4B3 (WT), and P2C1 (G983L).

### Growth factors and inhibitors

Recombinant human EGF (PeproTech AF-100-15), HGF (Thermo 100-39), IGF1 (PeproTech 100-11), and NRG1 (PeproTech 100-03) were used at concentrations of 10, 50, 10, and 37.5 ng/mL, respectively, unless otherwise noted. The EGFR inhibitor gefitinib (LC Laboratories G4408), MET inhibitor PHA-665752 (Santa Cruz Biotechnology sc-203186), PI3K inhibitor GDC-0941 (Cayman Chemicals 1160), JNK inhibitor SP60015 (LC Laboratories S-7979), and CREB inhibitor 666-15 (MedChemExpress HY-101120) were used at 10, 3, 1, 20, and 1 μM, respectively, unless otherwise noted. All inhibitors were reconstituted in DMSO according to manufacturer instructions. Prior to treatment with growth factors, cells were serum-starved overnight (∼18 hours) with in DMEM containing 0.1% fetal bovine serum.

### siRNA-mediated knockdown

For siRNA-mediated knockdowns, cells were plated and allowed to adhere overnight. Control siRNA (Cell Signaling Technology 6568) or HER3 siRNA (Cell Signaling technology 6422) were diluted in OptiMEM medium (Gibco 11058021), incubated with RNAiMAX lipofectamine (Thermo 13778030) for 5 minutes, and added to cells in complete medium for 72 hours, unless otherwise noted. 48 hours later, growth factors were added for 24 hours.

### Western blotting

HSC3 cells were lysed via scraping into a cell extraction buffer (Life Technologies FNN0011) supplemented with protease and phosphatase inhibitors and PMSF (Sigma-Aldrich 08340, P0044, and P5726). Crude lysates were clarified by centrifugation for 10 minutes at 20,800 rcf in a refrigerated benchtop centrifuge. Protein concentrations were measured using a bicinchoninic acid (BCA) assay (Pierce, Thermo 23225). To prepare samples for electrophoresis, 10-20 µg of protein was mixed with 4× NuPAGE LDS buffer (Pierce NP0007) and 500 mM dithiothreitol (Sigma D9779), and 20 µL was loaded per well onto 4-12% polyacrylamide gels (Invitrogen NP0336BOX). Electrophoresis was run, and gels were transferred to 0.2-µm nitrocellulose membranes (BioRad 1620168) using a BioRad TransBlot Turbo system. Blocking was done using 1:2 diluted Intercept Blocking Buffer (IBB) (Licor 927-60001). Primary antibodies were diluted in IBB and added at 4°C overnight on a rocker.

These antibodies were used for western blotting: GAPDH (Cell Signaling Technology 2118 and Santa Cruz Biotechnology sc32233), pAKT S473 (Cell Signaling Technology 9271), AKT (Cell Signaling Technology 9272), pERK 44/42 Thr 202/Tyr 204 (Cell Signaling Technology 4370), PTPRJ/DEP1 (Santa Cruz Biotechnology sc-376794), pSTAT3 Y705 (Cell Signaling Technology 9145), ERK (Cell Signaling Technology 4695 and 4696), pERK Thr 202/Tyr 204 (Cell Signaling Technology 4370), pMET Y1349 (Cell Signaling Technology 3133), pMET Y1234/1235 (Cell Signaling Technology 3126), HER3 (Cell Signaling Technology 12708), pHER3 Y1289 (Cell Signaling Technology 2842), EGFR (CST 4267), pEGFR Y1068 (CST 3777), MET (CST 8198), pMET Y1234/5 (CST 3077). Per manufacturer recommendations, all primary antibodies were diluted at 1:1000 in IBB prior to use, with the exceptions that total antibodies were diluted 1:2000 for Fig. 1 and GAPDH antibody was diluted 1:5000 for the rest of the figures. After primary incubations, membranes were washed once with 0.1% PBS-Tween for 5 min and twice with 1× PBS for 10 min before adding the species-appropriate IR-conjugated secondary antibodies for two hours at room temperature (1:10,000 or 1:20,000 anti-mouse 680 [Rockland #610-144-121] or anti-rabbit 800 [Rockland # 611-145-002]). Membranes were then washed once in 0.1% PBS-Tween for 5 min and twice with 1× PBS for 10 min before imaging on a LICOR Odyssey CLx.

**Figure 1.**
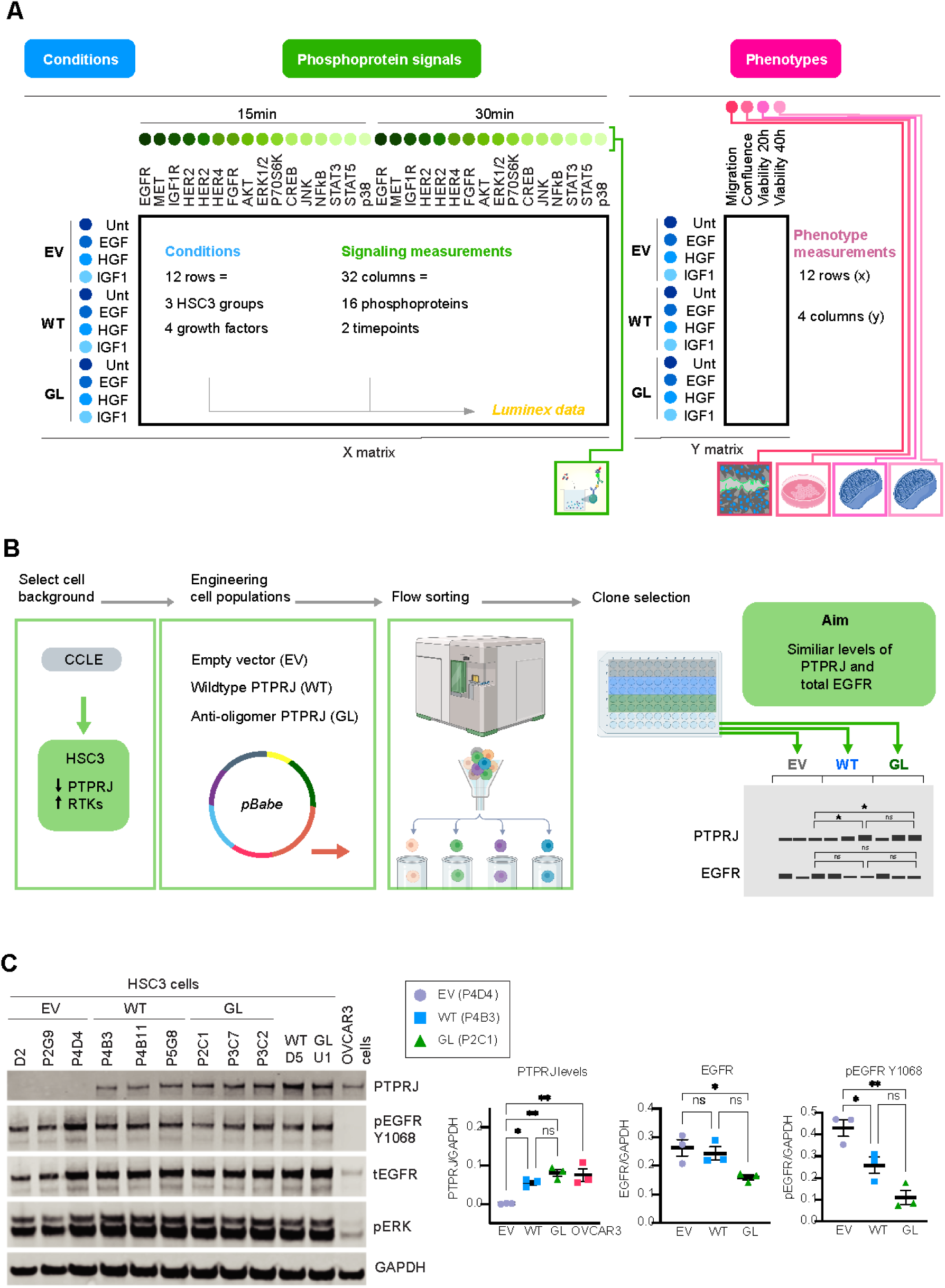
Experimental design and cell engineering. **(A)** Schematic depicting the HSC3 PTPRJ expression backgrounds, growth factor treatments, protein measurements at two time points, and four phenotypic measures. These data were used construct to matrices for partial least squares regression (PLSR). EV = empty vector; WT = wild type; GL = G983L. Created with BioRender. **(B)** Schematic depicting HSC3 cell population engineering, flow sorting, and selection of clones. Created with BioRender. **(C)** HSC3 cell clones were serum starved in 0.1% FBS DMEM media for 18 hours and treated with 10 ng/mL EGF for 15 minutes prior to lysis. Western blotting was performed for the indicated proteins. The clones selected for further experiments were P4D4, P4B3, and P2C1. Bars represent the mean ± SEM. * p < 0.05, ** p < 0.01, *** p < 0.001.

### Quantitative real-time PCR (qRT-PCR)

Cells were lysed from 6-well plates using RNAse Zap-treated cell scrapers in 500 µL solution of 1:50 Buffer RLT (QIAGEN #74104) at 2M DTT per well. Lysates were homogenized using QIAshredder spin columns (QIAGEN #79654). RNA extraction and purification was done using an RNA extraction kit, following manufacturer instructions for the lysis of adherent mammalian cells (QIAGEN RNeasy kit #74104), including an optional gDNA elimination step (QIAGEN RNase-free DNAse Set #79254). RNA quality and concentration was read using a spectrophotometer (NanoDrop 2000c/2000 ThermoFisher). RNA was diluted to a concentration of 5 μg in 10 μL nuclease-free water (unless otherwise specified). Reverse transcription to cDNA was done using the High-Capacity cDNA Reverse Transcriptase Kit (Applied Biosciences # 4368814). PowerUp SYBR Green (Applied Biosciences: #4367659) detection was used for qRT-PCR. We plated three biological replicates and measured two technical replicates in 96 well plates, unless otherwise specified. Relative changes in transcript abundances were determined using the 2^−ΔΔCT^ method. All transcripts were normalized to *GAPDH*. Primers were designed using NCBI PrimerBlast or primer3 and were designed to span two adjacent exons to avoid detecting genomic DNA. Alternatively, published or lab-validated primers were used. Products of primers not previously validated were run on a gel to confirm the bands had the expected molecular weights and to check for the formation of dimers. Primer sequences are given in Supplemental Table 1.

### Live-cell nucleus counts

HSC3 cells (EV, WT, G983L; n=4 per condition) were plated at 2,500 cells/well in 96-well plates with 100 µL complete medium per well. The next day, control siRNA or HER3 siRNA plus RNAiMAX lipofectamine diluted in OptiMEM were added according to manufacturer instructions and left on for ∼72 hours before imaging. 24 hours prior to imaging, cells were treated with or without 50 ng/mL HGF, diluted in serum-free OptiMEM. On the day of imaging, NucBlue (Thermo R37605) was added to the cells according to manufacturer instructions, and wells were imaged on a BioTek Cytation 5 at 10× magnification one hour later. Nuclei were counted in the DAPI channel using the Cytation *Cellular Analysis* program for cell counts. Six images at different locations were taken in each well. Plates were later fixed and imaged after nuclei counting following the immunofluorescence protocol.

### Immunofluorescence microscopy

HSC3 cells in 96-well plates were washed twice with ice-cold PBS, fixed using 50 µL 4% paraformaldehyde in PBS for 20 minutes, washed twice with PBS, then permeabilized using 0.25% Triton-X-100 in PBS at room temperature for 5 minutes. After washing twice with 1× room-temperature PBS using the BioTek 406 EL Washer-Dispenser, 35 µL of primary antibodies in IBB was added per well, and the plate was incubated overnight in a 4°C humidified chamber. The complete list of primary and secondary antibodies used can be found in Supplemental Table 2. The next day, wells were washed with PBS using the plate washer and 35 µL of 1:750 secondary antibodies and 1:2,000 DNA stain Hoechst in IBB were added for one hour at room temperature. Wells were washed with PBS, then all buffer was aspirated and 50 µL 20% filtered glycerol in PBS was added as imaging buffer. The BioTek Cytation 5 was used to image immunofluorescence on the CY5, RFP, and DAPI channels.

### Automated image analysis for fluorescence microscopy

For each channel, the LED brightness, gain, and exposure time were calibrated to the well with the highest intensity as imaged via manual mode. If there were different antibodies (e.g., EGFR and MET) on different wells of the same plate that could be read on the same channel (e.g., RFP), these were calibrated separately. Six non-overlapping images were taken per well using a 10× objective. Cytation 5 rolling-ball background subtraction was implemented. To reduce blurriness, deconvolution was done via the Cytation 5 automated Richardson-Lucy deconvolution recommended per objective size. To quantify images, CellProfiler was used to measure intensities within individual cells. Nuclei were used as primary objects, and EGFR, HER3, or ERK were used as secondary objects for cell segmentation and identification. The integrated intensity and median intensity fluorescence of the desired measurement was taken of each cell object, and the integrated intensity used for further analyses.

### Scratch assays

HSC3 cells were seeded at 200,000 cells per well in a 12-well plate and grown for 48 hours in complete DMEM to form a confluent monolayer prior to overnight serum starvation. Cells were scratched with a 200-µL pipette tip, and excess media with cellular debris was removed, leaving 1 mL of media to cover the cells. Cells were then treated with vehicle or growth factors and monitored as wounds closed. Four images per well were taken every 2 hours for 24 hours, or until the wound closed, using a Curosis Celloger Mini Plus live-cell imaging system at 5% CO_2_ and 37°C. Percent closure was calculated and reported as previously described in (29, 30) using the ImageJ MRI Wound Healing Tool.

### xCELLigence measurements

The xCELLigence RTCA dual purpose (DP) system (Agilent) was used for real-time cell analysis (RTCA) experiments. As cells adhere and grow on the surface of the electrodes, changes in impedance for each well are recorded and reported as cell index values. The measurements reflect changes in cell adherence, morphology, and viability. Prior to seeding cells, the xCELLigence software was calibrated via a blank scan of cell culture medium to provide a baseline measurement. For each well of the RTCA E-plate (Agilent #6465412001), 50 µL of media was deposited to make a background measurement. 5000 cells were plated into each well to a final volume of 100 µL in complete DMEM. Growth factors were diluted in separate stocks to double the working concentration in complete DMEM. 100 µL of each growth factor-containing solution or media alone (control) was added to appropriate wells. Cells were allowed to equilibrate and settle in a cell culture hood at room temperature for 30 minutes prior to loading the plate onto the xCELLigence instrument. Measurements were taken every 15 minutes for 250 scans total (∼62 hours). For each condition, we normalized the change in cell index to the time point where initial adherence had settled and reported the slope of cell index versus time for 5 to 20 hours (the linear growth range for this cell type).

### MTT assay

The MTT (3-(4,5-dimethyl-2-thiazolyl)-2,5-diphenyl-2H-tetrazolium bromide) assay was used to measure cell viability and mitochondrial activity. 5,000 HSC3 cells in 50 µL complete media were plated per well in a 96-well plate. An additional 50 µL of growth factors in complete media were added to each well so that the final concentrations were 10 ng/mL EGF, 50 ng/mL HGF, or 10 ng/mL IGF1 (100 µL/well final volume). For control wells, another 50 µL of complete media was added. Cells were then grown for 20 or 40 hours. At the end of an experiment, 10 µL of 5 mg/mL MTT reagent in PBS was added per well, and plates were incubated for 2 hours at 37°C. Media was removed, and the remaining MTT crystals were solubilized in 200 µL DMSO. Plates were shaken to dissolve crystals, and absorbance at 580 nm (10 nm bandwidth) was measured using a plate reader. Each set of HSC3 clones was normalized to samples without growth factor and was reported as a percent viability in comparison to the control (normalized to 100% viability).

### Luminex assays

HSC3 cells were serum starved overnight in 0.1% FBS in DMEM. They were then treated for 15 or 30 minutes with EGF 10 ng/mL, HGF 50 ng/mL, IGF1 10 ng/mL, or vehicle. 12 conditions (3 PTPRJ expression backgrounds treated with 4 different growth factor conditions) were measured at the two time points, each with three biological replicates. Cells were immediately lysed after growth factor treatment, and protein concentrations were measured by BCA assay. For Luminex assay plates, 25 µL of total volume was loaded per well, which contained 5 or 10 µg of total protein for the Multipathway or RTK kit, respectively. Luminex data were analyzed using the xPONENT program and xMAP INTELLIFLEX® High Sensitivity operating mode on a Luminex MAGPIX system at the University of Virginia Flow Cytometry Core Facility. We used the background-subtracted mean fluorescence intensity (MFI) for analysis. Some IGF1R measurements show as negative due to the background being higher than analyte concentrations. Multiplex MILLIPLEX Cell Signaling kits were used according to manufacturer instructions. We used the Milliplex^®^ MAP 9-plex Multi-Pathway Signaling Magnetic Bead kit (48-680MAG) to probe for ERK/MAP kinase 1/2 (Thr 185/Tyr 187), Akt (Ser 473), STAT3 (Ser 727), JNK (Thr 183/Tyr 185), p70S6K (Thr412), NFkB (Ser 536), STAT5A/B (Tyr 694/699), CREB (Ser 133), and p38 (Thr 180/Tyr 182). We used a custom Milliplex^®^ MAP 7-plex Human RTK Mitogenesis Total Protein Magnetic Bead kit (HPRTKMAG-01-07) to probe total pan-tyrosine phosphorylation of EGFR, MET, IGF1R, HER2, HER3, HER4, and FGFR. Analytes with high signal-to-noise or poor linearity were excluded from the analysis.

### Principal component analysis

PCA was performed on multiplexed data in R.

### Partial least squares regression

Partial least squares regression (PLSR) was performed in MATLAB using the built-in *plsregress* function. We first developed four different PLSR models, one for each phenotypic outcome, including cell migration via scratch assay, xCelligence cell index, and MTT assay at 20 hr or 40 hr. For each model, the independent variable matrix consisted of the mean-centered and variance-scaled Luminex measurements from EV, WT, and G983L cells treated with EGF, HGF, IGF1 or vehicle for 15 or 30 min. These measurements formed a 12 × 11 independent variable matrix, where rows represented experimental conditions and columns represented signaling features at each treatment time point. The corresponding functional assay measurement was used as the dependent variable, forming a 12 × 1 dependent variable matrix (or vector). We also developed a PLSR model in which all four phenotype vectors were concatenated to form a single 12 × 4 dependent variable matrix. Model performance was evaluated using leave-one-out cross validation, in which one observation was withheld as the test set and the remaining observations were used for model training. PLSR models were evaluated based on goodness of fit (R^2^Y) and predictive power (Q^2^Y) based on cross-validation. To identify signaling features that contributed strongly to variation in the response variable, we calculated variable importance in projection (VIP) scores. VIP scores reflect the contribution of each feature across all retained PLS components, computed by weighting a feature’s contribution to each PLS component by the amount of variance in the response variable explained by that PLS component. VIP-enriched models were created by regenerating PLSR models after eliminat features with VIP scores less than 1. To visualize relationships among treatments, cell types, signaling features, and functional outcomes, we generated biplots using the scores and loadings from the first two PLS components. These components captured the majority (> 50%) of variance in the dependent variable.

### Statistical analysis

For all experiments where statistics were computed, at least three biological replicates were analyzed. Statistical analyses were performed using Prism Graphpad. For Luminex assays, analysis was done via two-way ANOVA for the PTPRJ factor, GF treatment factor, and interaction factor with Tukey’s multiple comparisons post-hoc test. Scratch, xCelligence and MTT analyses were conducted using a one-way ANOVA with Sidak’s multiple comparisons test. For PCR, fold changes were analyzed via one-way ANOVA with Dunnett’s test, for comparisons involving three or more groups. Western blot data were reported as the signal intensity (with median background subtraction) divided over loading control (e.g., GAPDH or beta-actin). For tests of two groups, Welch’s *t* test was conducted. For tests of three or more groups, one-way ANOVA was performed, with Sidak’s multiple comparisons post hoc testing.

## RESULTS

### PTPRJ expression background and growth factor context contribute to signaling variance

To generate a dataset capturing PTPRJ regulation of signaling networks and phenotypes, HSC3 cells engineered with ectopic expression of WT or G983L PTPRJ, or an empty vector-transduced control, were treated with EGF, HGF, or IGF1. HSC3 cells were chosen due to their low endogenous PTPRJ expression, abundant EGFR, MET, and IGF1R expression, and responsiveness to EGFR signaling (29, 41, 42). Cells were treated for 15 or 30 min with growth factors, and lysates were gathered for multiplexed phosphoprotein measurements. Four phenotypic measurements were made after growth factor treatments. The schematic in Figure 1A illustrates how the resulting data were used to form matrices of independent and dependent data to generate PSLR models (Fig. 1A). The clonal cell lines used were isolated and evaluated by immunoblotting as described in *Methods* (Fig. 1B). Clones with similar EGFR expression and similar PTPRJ expression (for WT and G983L) were selected for experiments (Fig. 1C).

As expected, EGF, HGF, and IGF1 increased the phosphorylation of EGFR, MET, and IGF1R, respectively (Fig. S1). The phosphorylation of those receptors was reduced by PTPRJ expression, with the largest effect observed for the G983L mutant. PTPRJ wildtype and mutant also significantly suppressed HER3 phosphorylation for all growth factor conditions and had some effects on HER2 phosphorylation. To our knowledge, regulation of HER3 tyrosine phosphorylation by PTPRJ has not previously been reported. PTPRJ expression also altered the phosphorylation of some of the downstream effectors measured, with effects observed for multiple growth factor settings for CREB and JNK.

The mean-centered and variance-scaled measurements are shown in Fig. 2A as a 12 × 32 matrix of signaling data and a 12 × 4 matrix of phenotype data. To understand the major sources of signaling variance, we used principal component analysis (PCA) to reduce the dimensionality. Principal components 1 and 2 (PC1 and PC2) explained 41% and 19% of the variance in the signaling data, respectively (Fig. 2B). Grouping measurements in the PCA projection demonstrated that differences in PTPRJ expression and growth factors contributed roughly equally to variance in the data. HGF, EGF, and IGF1 conditions projected toward the upper right, lower right, and left/upper left, respectively. Within groups of a common ligand and treatment time, variance was sometimes relatively small (e.g., IGF1 15 min) or large (HGF or EGF 15 min). The variance observed within groups was in some cases as large as the variance across groups (e.g., from HGF 30 min to EGF 15 min). In terms of feature loadings, measurements of STAT3, ERK1/2, CREB, and p38 projected strongly in both PCs, STAT5, HER2, EGFR projected strongly in PC1, and AKT and p70S6K projected highly in PC2 (Fig. 2B). A scree plot shows that a third PC would have added only an additional 10% explanation of the variance (Fig. 2C).

**Figure 2.**
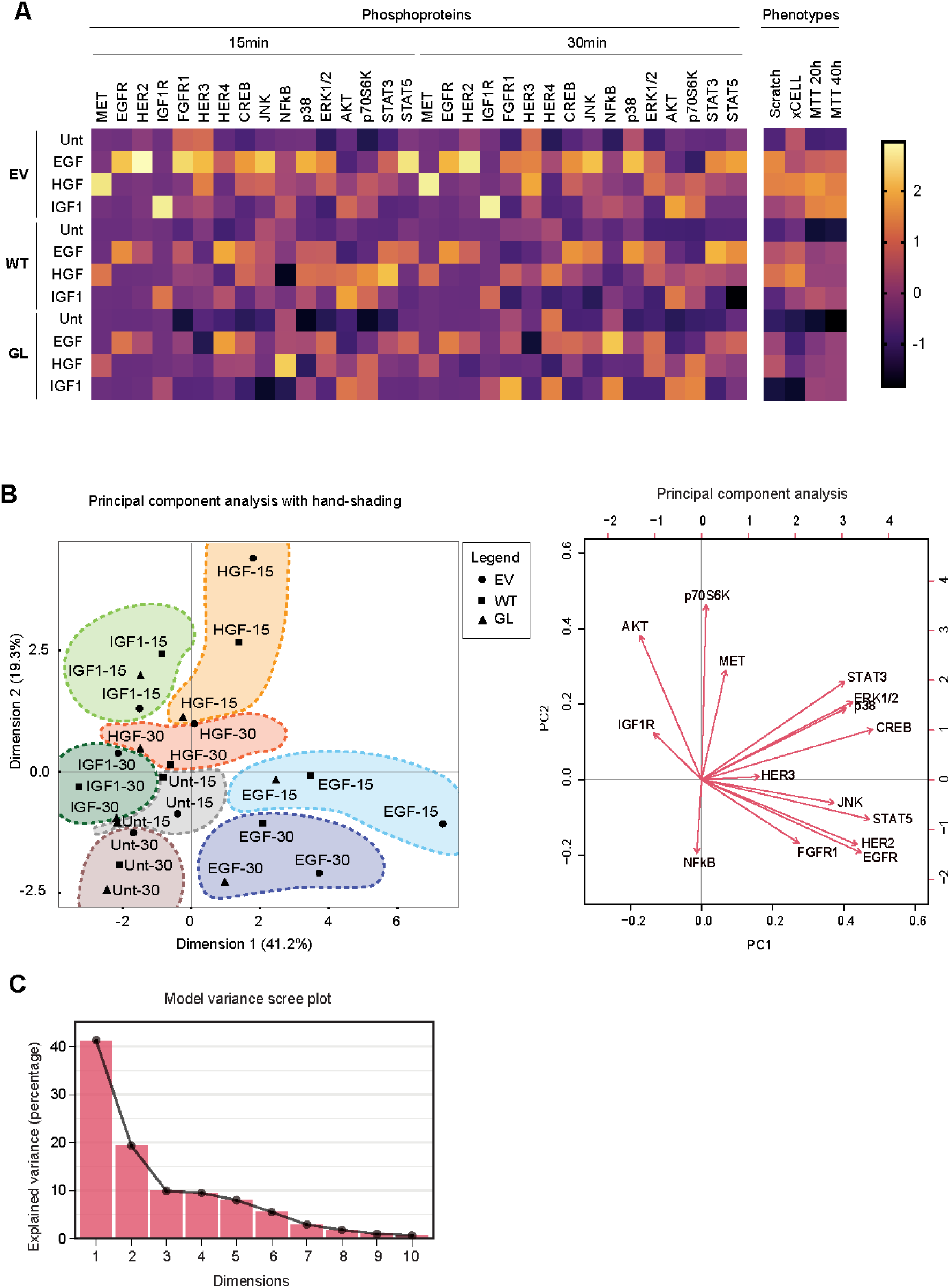
PTPRJ expression and growth factor conditions provide contribute similar variance to signaling measurements. **(A)** HSC3 cells were serum-starved for 18 hours prior to treatement with growth factors (10 ng/mL EGF, MET 50 ng/mL, IGF1 10 ng/mL, or untreated) for fifteen or thirty minutes. Lysates were used to run Luminex assays for the analytes indicated in columns. The heatmap shows Luminex data after mean centering and variance scaling for each column (i.e., for each analyte at each time point). EV = empty vector; WT = wild type; GL = G983L. Also shown are the scaled measurements for four phenotypes. **(B)** Principal component analysis was performed on the Luminex data shown in (A). The plot at left is hand-shaded to group clusters of the three PTPRJ expression settings by similar growth factor and treatment time. The biplot at right shows the projection of the features in the same principal component space. **(C)** Scree plot of the percentage explained variance per dimension. Bars represent the mean ± SEM. * p < 0.05, ** p < 0.01, *** p < 0.001.

### PTPRJ suppresses growth factor-driven cell migration

To identify signaling proteins that controlled the phenotypic outcomes we measured, we used PLSR. Like PCA, PLSR method effects a dimensionality reduction, but the latent variables (LVs, combinations of measured features) are selected to maximally capture variance in the independent and dependent variables simultaneously. We began with a model of the wound healing phenotype alone. Growth factors, especially EGF and HGF, promoted faster wound closure than control conditions, but this effect was suppressed by PTPRJ, especially the G983L mutant (Fig 3A,B). In a PLSR model with two latent variables, ERK, JNK, CREB, and HER3 co-projected with the wound healing (scratch) phenotype (Fig. 3C). The predicted importance of HER3 was removed, however, after VIP enrichment of the model. HGF was the strongest driver of the signals that most influenced scratch closure (Fig. 3D). The model also captured the preferential ability of G983L PTPRJ to antagonize wound healing in the context of IGF1. Metrics of quality for the unenriched and VIP-enriched PLSR models and VIP scores by feature for both models are shown in Figs. 3E and 3F, respectively.

**Figure 3.**
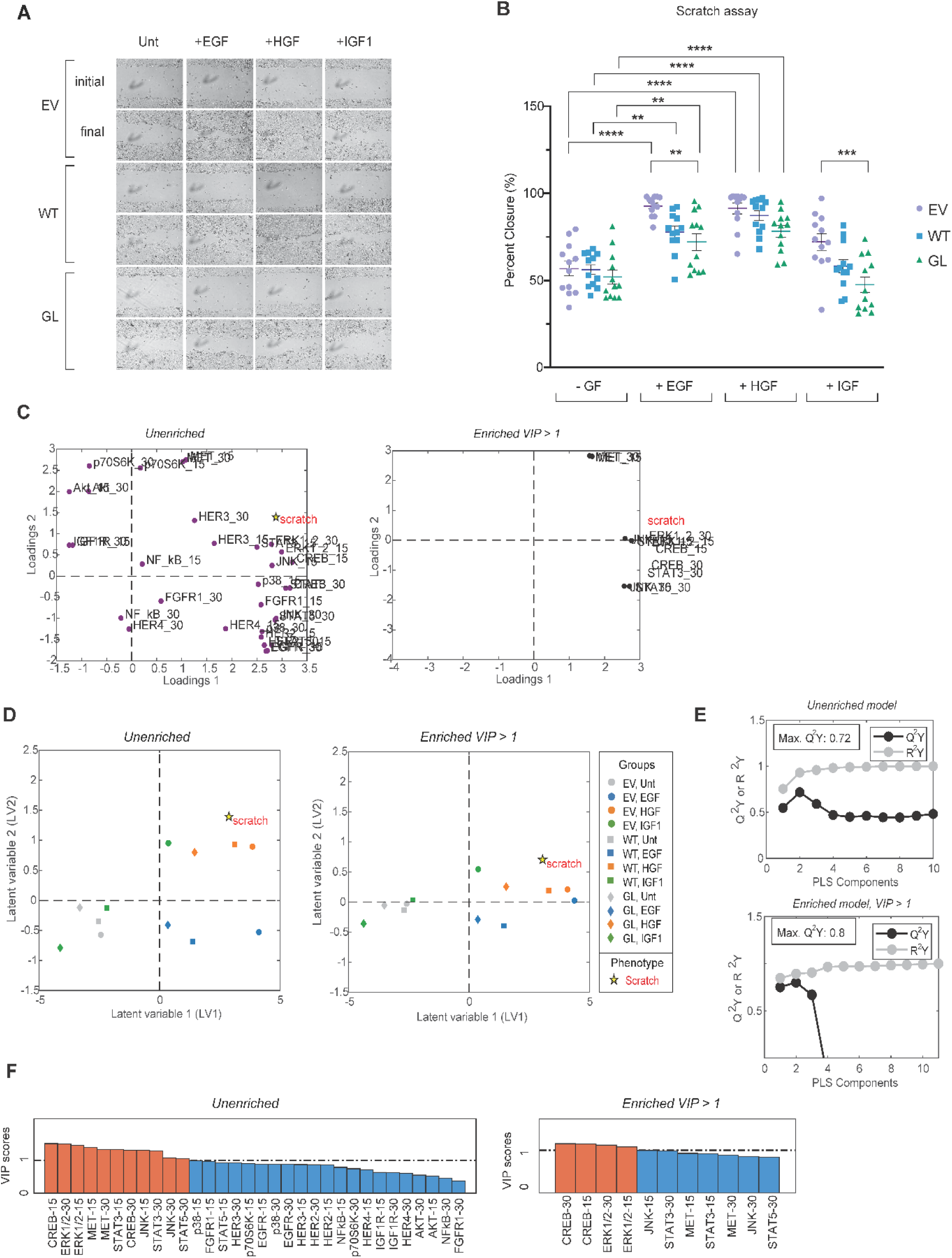
PTPRJ G983L impedes cell migration driven by multiple growth factors. **(A)**HSC3 cells were serum-starved for 18 hours before treatment with growth factors. Representative phase-contrast images of the scratch area are shown at the beginning and end of the experiment. EV = empty vector; WT = wild type; GL = G983L. **(B)** Percent wound closure was quantified from the images. Quantification of PTPRJ WT and GL percent wound closure compared to EV cells. One-way ANOVA with Sidak’s multiple comparisons test; * p < 0.05, ** p < 0.01, *** p < 0.001, ns = not significant. **(C)** A PLSR model was generated based on Luminex data as the independent variables and scratch assay data shown in (B) as the dependent variable. Loadings are shown in biplots of the first two latent variables for models before or after VIP enrichment. **(D)** Scores plots for the 12 experimental conditions are shown before and after VIP enrichment. **(E)** Model R^2^Y and Q^2^Y coefficients are shown as a function of the number of PLS components (or latent variables). **(F)** VIP scores for the signaling measurements are shown for the unenriched or VIP-enriched models.

### PLSR model of xCELLigence cell index

The second phenotype measured was the xCELLigence-based cell index, an impedance-based metric reflecting the aggregate effects of cell confluence, adhesion, and morphology. For all growth factor conditions, the rate of change of cell index was most suppressed in the G983L background (Fig 4A,B). HER3 and MET loadings co-projected most strongly with the change in cell index (xCELL) in the unenriched PLSR model, and HER3, JNK, STAT3, and NF-kB co-projected with xCELL along LV1 in the enriched model (Fig 4C). Scores plots reflected the strong negative regulation of cell index changes by PTPRJ G983L and the ability of HGF to promote it most strongly (Fig. 4D). The unenriched PLSR model had a maximum Q^2^Y = 0.49 for 10 latent variables, and the VIP-enriched model had a stronger Q^2^Y = 0.75 for seven latent variables (Fig. 4E). Of the features with VIP > 1 for the enriched model, HER3, JNK and STAT3 had relatively large LV1 loadings, whereas NFκB had a larger LV2 loading. Comparing the enriched weights and scores plots in Fig. 4C and D, one could conclude that the PTPRJ mutant decreased the rate of change of the cell index either by promoting NFkB signaling or by suppressing signaling involving JNK, CREB, and STAT3. Overall, WT PTPRJ did not exert strong control over the cell index phenotype, although it did significantly control some signaling pathways. Thus, the PLSR model identified the pathways preferentially controlled by the PTPRJ mutant and most regulate cell index changes.

**Figure 4.**
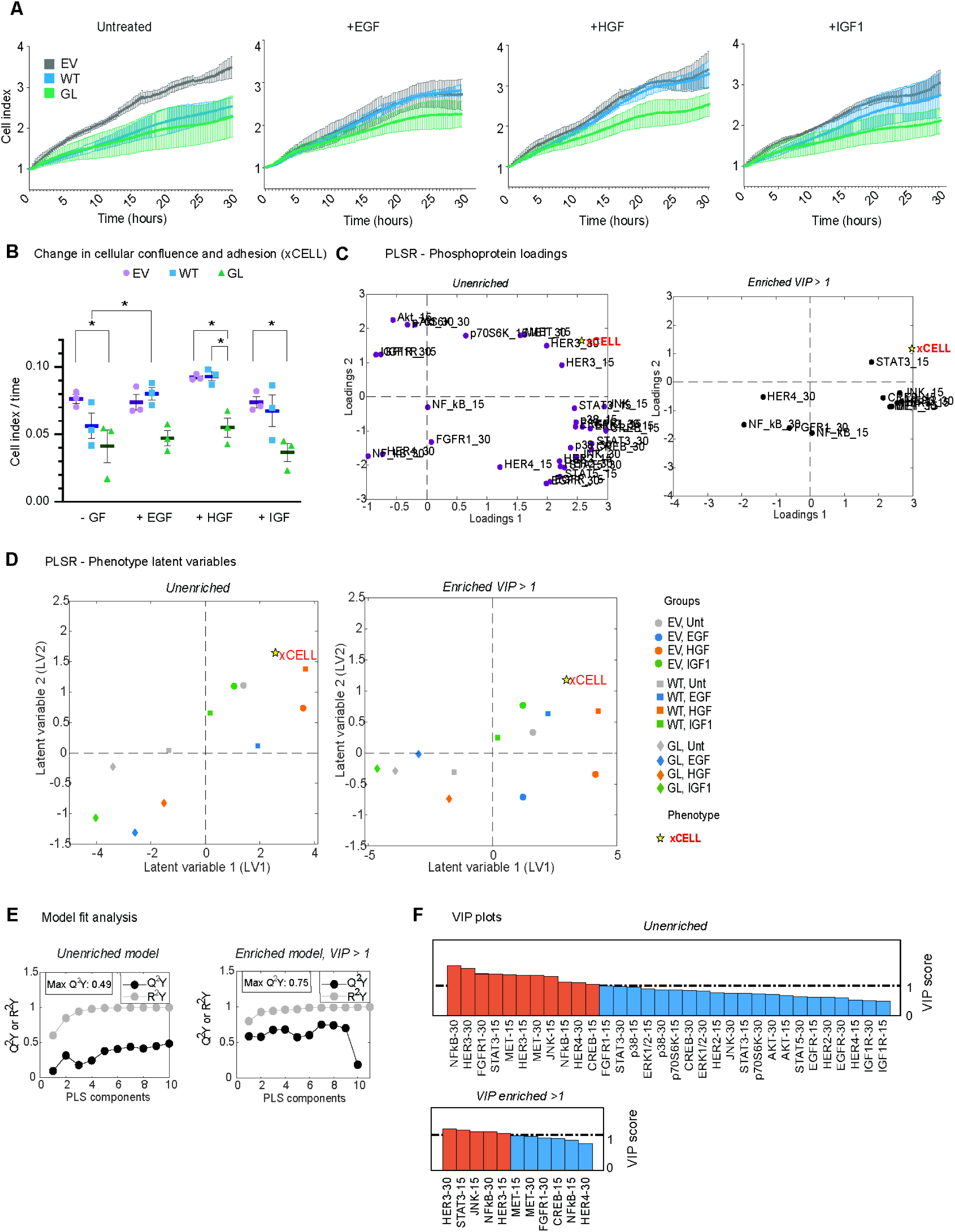
PLSR modeling nominates HER3, pCREB, and NFκB as PTPRJ-regulated signaling nodes that regulate cell index. **(A)** HSC3 cells were treated with growth factors (EGF 10 ng/mL, HGF 50 ng/mL, IGF1 10 ng/mL) in serum-free medium and allowed to adhere to xCell plates between 0-5 hr. The cell index was measured by xCELLigence RTCA in 15-minute increments for 0 – 30 hours for the indicated cell lines and conditions. The cell index was normalized to compensate for initial adhesion. EV = empty vector; WT = wild type; GL = G983L. **(B)** The data shown in panel (A) were used to compute rates of change of cell index over time from 5 - 20 hr. One-way ANOVA with Sidak’s multiple comparisons test; * p < 0.05, ** p < 0.01, ns = not significant. Bars represent mean ± SEM. **(C)** A PLSR model was created based on Luminex data as independent variables and the cell index data in panel (B) as the dependent variable. Loadings are shown in biplots of the first two latent variables for models before or after VIP enrichment. **(D)** Scores plots for the 12 experimental conditions are shown before and after VIP enrichment. **(E)** Model R^2^Y and Q^2^Y coefficients are shown as a function of the number of PLS components (or latent variables). **(F)** VIP scores for the signaling measurements are shown for the unenriched or VIP-enriched models.

### PLSR models of cell viability assays

Continuing with the development of models for individual phenotypes, we next performed PLSR for the MTT assays at 20 or 40 hours. PTPRJ WT and G983L expression typically led to equivalent decreases in MTT signal across growth factor conditions at 20 hours (Fig. 5A). In the unenriched two-component PLSR model, p70S6K, MET, and HER3 co-projected most strongly with MTT signal (Fig. 5B). In the VIP-enriched version of the model, MET was the strongest co-projecting feature. Scores plots for the unenriched and VIP-enriched models demonstrated the strong dependence of viability on growth factor treatments and the tendency for all growth factors to promote viability especially for the EV setting (Fig. 5C). Performance of the unenriched model was good for three of more LVs, and this was improved with VIP enrichment but with a non-monotonic effect with increasing LVs (Fig. 5D). AKT-related signaling nodes were prominently represented among the top features in both models (Fig. 5E). Overall, similar trends were observed for MTT measurements and models at 40 hours (Fig. 6), with the exception that JNK arose as an important phenotype-controlling feature.

**Figure 5.**
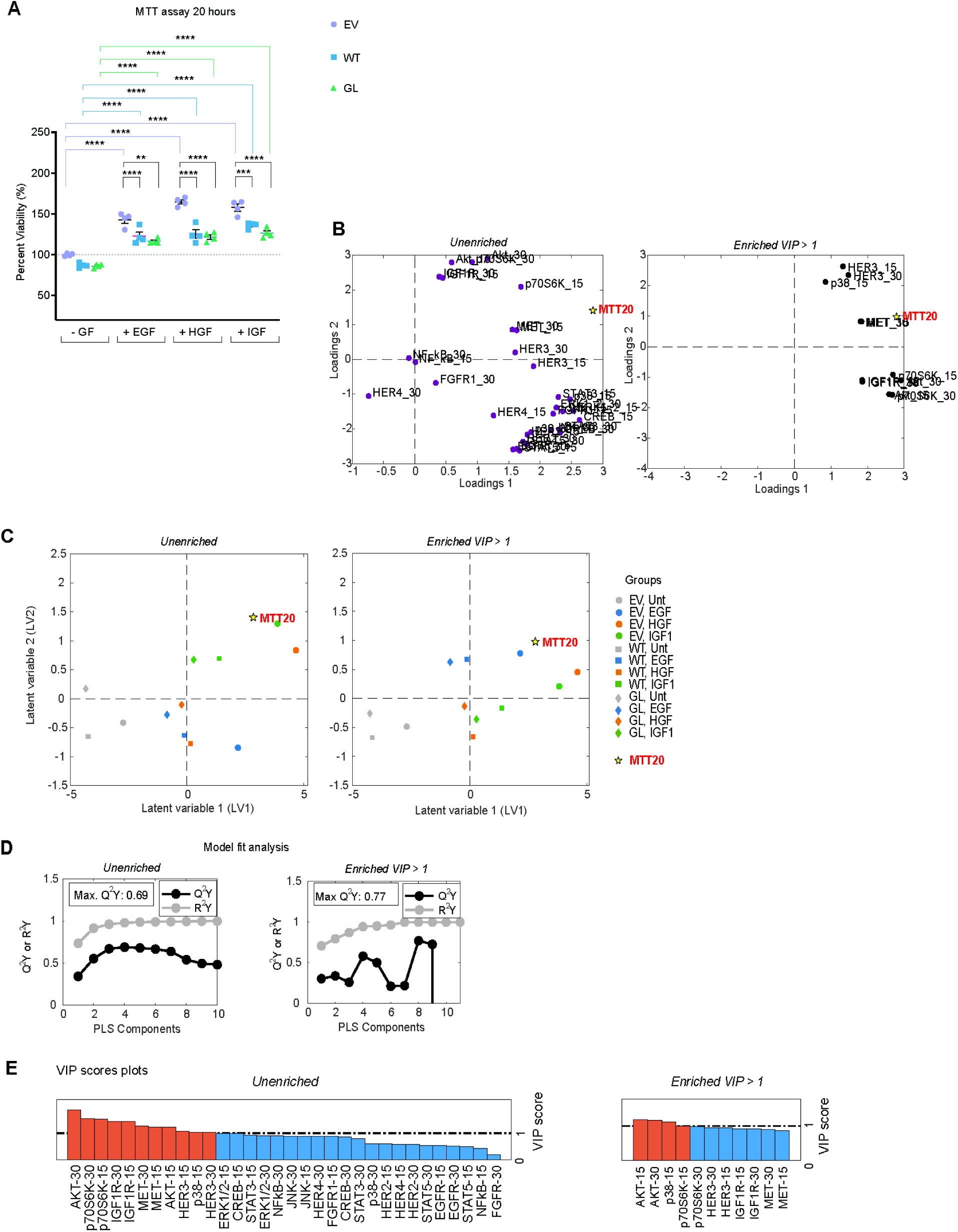
PTPRJ reduces growth factor-induced changes in cell viability through PLSR-identified signaling pathways (20-hour MTT assay). **(A)** Cells were treated with 10 ng/mL EGF, 50 ng/mL HGF, or 10 ng/mL IGF1 growth factor and grown for 20 hours before conducting an MTT assay. Percent viability is reported as a comparison against the untreated empty vector (EV), which was normalized to 100% viability. EV = empty vector; WT = wild type; GL = G983L. Bars represent the mean ± SEM. EV = empty vector; WT = wild type; GL = G983L.One-way ANOVA with Sidak’s multiple comparisons test; * p < 0.05, ** p < 0.01, *** p < 0.001, **** p < 0.0001. **(B)** A PLSR model was created using the Luminex data as independent variables and the phenotype data shown in (A) as the dependent variable. Loadings are shown in biplots of the first two latent variables for models before or after VIP enrichment. **(C)** Scores plots for the 12 experimental conditions are shown before and after VIP enrichment. **(D)** Model R^2^Y and Q^2^Y coefficients are shown as a function of the number of PLS components (or latent variables). **(E)** VIP scores for the signaling measurements are shown for the unenriched or VIP-enriched models.

**Figure 6.**
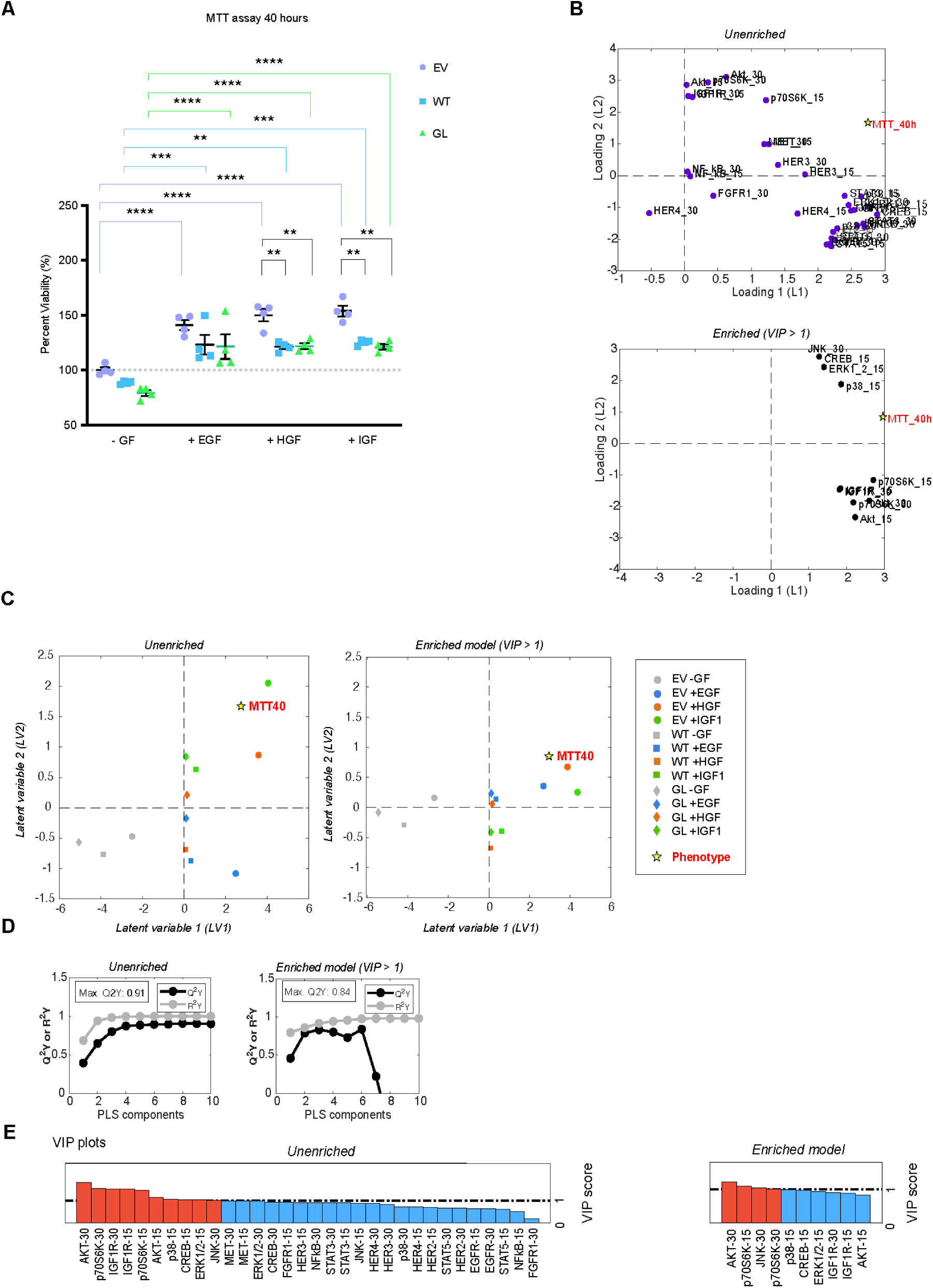
PTPRJ reduces growth factor-induced changes in cell viability through PLSR-identified signaling pathways (40-hour MTT assay). **(A)** Cells were treated with 10 ng/mL EGF, 50 ng/mL HGF, or 10 ng/mL IGF1 growth factor and grown for 40 hours before conducting an MTT assay. Percent viability is reported as a comparison against the untreated empty vector (EV), which was normalized to 100% viability. EV = empty vector; WT = wild type; GL = G983L. Bars represent the mean ± SEM. One-way ANOVA with Sidak’s multiple comparisons test; * p < 0.05, ** p < 0.01, *** p < 0.001, **** p < 0.0001. **(B)** A PLSR model was created using the Luminex data as independent variables and the phenotype data shown in (A) as the dependent variable. Loadings are shown in biplots of the first two latent variables for models before or after VIP enrichment. **(C)** Scores plots for the 12 experimental conditions are shown before and after VIP enrichment. **(D)** Model R^2^Y and Q^2^Y coefficients are shown as a function of the number of PLS components (or latent variables). **(E)** VIP scores for the signaling measurements are shown for the unenriched or VIP-enriched models.

### A PLSR model based on all phenotypes nominates CREB, AKT, STAT3, ERK1/2, and JNK

To gain understanding of network level control of all phenotypes simultaneously, we created a PLSR model including the four phenotype measurements together (Fig. 7). In both the unenriched and VIP-enriched models, the scratch and xCELL phenotypes co-projected more closely than they did with the MTT phenotype (Fig. 7A). This may reflect a heavy component of cell migration in determining the cell index reported by the xCELLigence platform. Both phenotypes were explained by similar signals, in that all phenotypes projected equally and in the same direction in the dominant first latent variable. The main difference between the two phenotype groupings thus arose in LV2, and feature loadings reflected a heavier dependence of MTT signals on AKT pathway components and of scratch and xCELL signals on HER3, STAT3, CREB, and JNK. Scores plots reflected a tendency for EGF and HGF to drive the scratch and xCELL phenotypes preferentially, and IGF1 to drive the MTT phenotypes (Fig. 7B). Both the unenriched and VIP-enriched PLSR models met standard quality thresholds, but, unlike the models of individual phenotypes, model quality did not improve overall with VIP enrichment (Fig. 7C). CREB, AKT, STAT3, ERK, and JNK were common features with VIP > 1 for the unenriched and VIP-enriched models, consistent with aggregate inferences from prior models based on individual phenotypes (Fig. 7D).

**Figure 7.**
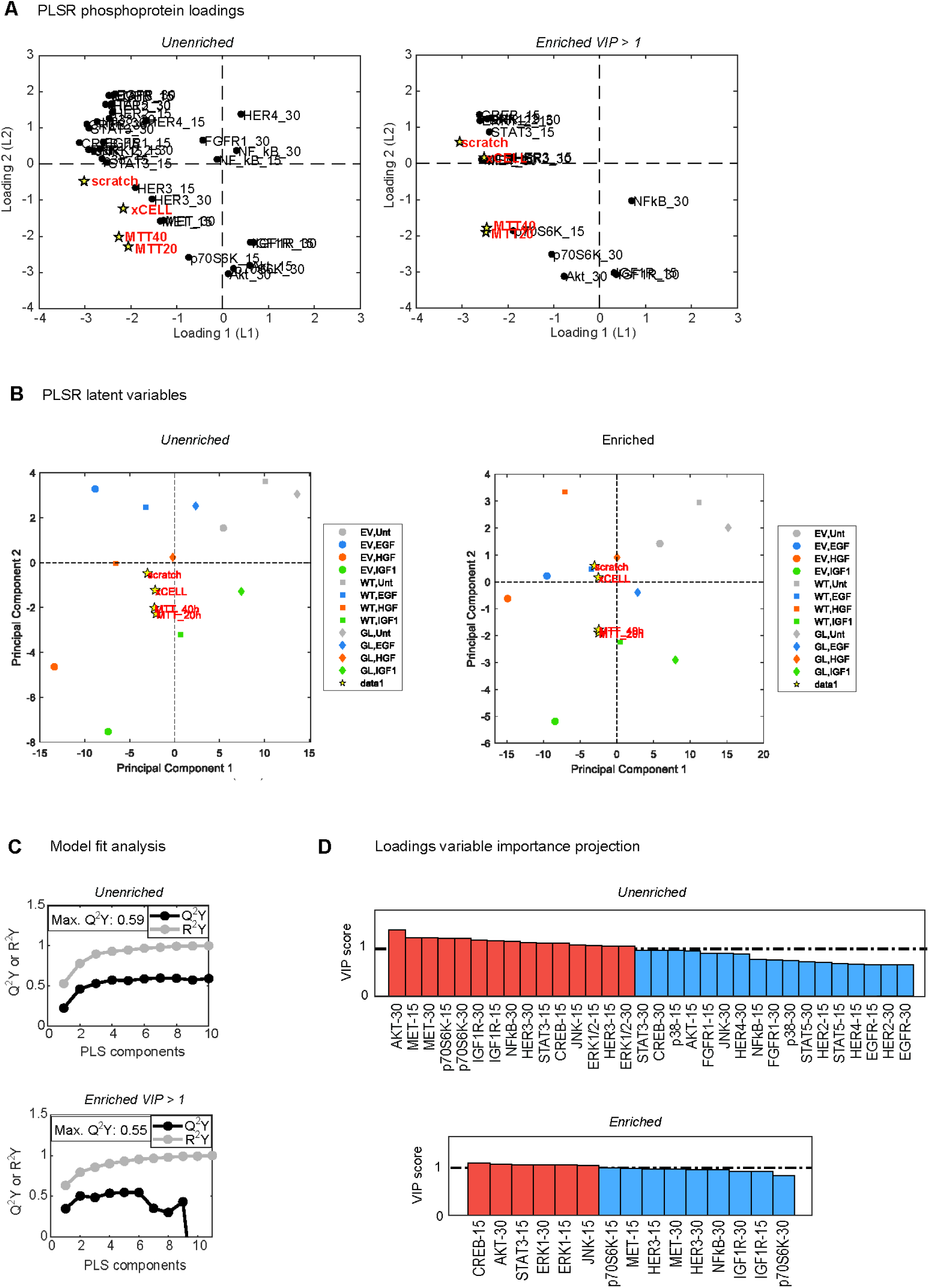
A PLSR model based all phenotypes groups PTPRJ-regulated phenotypes and the pathways that most control them. **(A)** A PLSR model using the Luminex data as independent variables and all four phenotype measurements as dependent variables. Loadings are shown in biplots of the first two latent variables for models before or after VIP enrichment. **(B)** Scores plots for the 12 experimental conditions are shown before and after VIP enrichment. **(C)** Model R^2^Y and Q^2^Y coefficients are shown as a function of the number of PLS components (or latent variables). **(D)** VIP scores for the signaling measurements are shown for the unenriched or VIP-enriched models.

### Role of HER3 in PTPRJ-regulated signaling and phenotypes

Based on the frequently predicted importance of HER3 in PLSR models, we undertook experiments to investigate the role of HER3 in regulating HSC3 cell signaling and phenotypes as a function of PTPRJ expression. HER3 is a pseudokinase and ErbB family member that forms dimers or trimers with EGFR, HER2 and HER4 to activate downstream signaling (43, 44). We began by testing the effect of siRNA-mediated HER3 depletion on cell proliferation with or without exogenous HGF, given the model-predicted importance of that ligand condition for PTPRJ-regulated effects (Fig. 8A). Without HGF, there was no apparent difference in cell proliferation among EV, WT, and G983L cells for the control siRNA, but WT and G983L cells exhibited a modest reduction in proliferation compared to EV cells (less than two-fold) when HER3 was depleted. In the presence of HGF, WT and G983L cells exhibited a more than two-fold decrease in proliferation compared to EV for the control siRNA condition, and HER3 depletion brought cell counts for all three cell lines to a similar level. We interpreted these results collectively to indicate that HER3 promoted proliferation in the presence of HGF through a mechanism that can be attenuated by WT or mutant PTPRJ.

**Figure 8.**
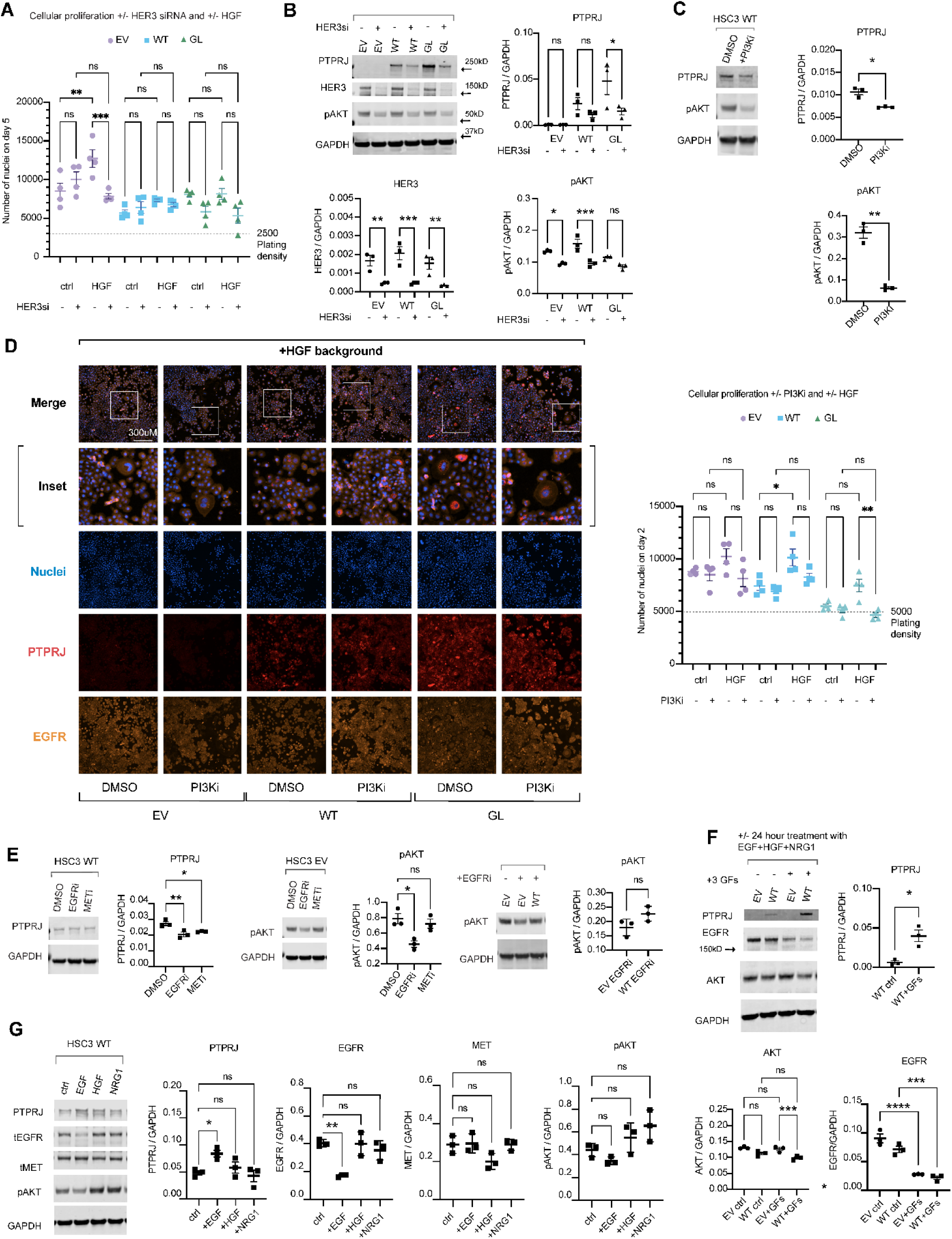
PTPRJ participates in an unanticipated feedback with HER3, RTKs, and AKT. **(A)** HSC3 cells were transfected with control or HER3 siRNA for a total of 72 hours. 50 ng/mL HGF or vehicle was added 24 hours before imaging. Live-cell DAPI imaging and nucleus counting was performed (n = 4 wells per group). EV = empty vector; WT = wild type; GL = G983L. One way ANOVA and post hoc Sidak’s multiple comparisons test. **(B)** HSC3 cells were transfected with control or HER3 siRNA for 48 hours, and 50 ng/mL HGF was added to each for 15 minutes prior to cell lysis (n = 3 per group). Western blotting was conducted for the indicated proteins (second gel with third set of replicates not shown). **(C)** HSC3 WT PTPRJ cells were treated with DMSO or 1 µM GDC-0941 for 3 days, and 50 ng/mL HGF was added 15 minutes prior to lysis. Western blotting was conducted for the indicated proteins. Welch’s t-test. **(D)** HSC3 cells were treated with DMSO or 1 µM GDC-0941 for 3 days, and 50 ng/mL HGF or vehicle was added to appropriate conditions 24 hours before nucleus counting and DAPI imaging (n = 4 per group). Cells were fixed, and immunofluorescence imaging was performed for the indicated targets (n = 1 for EGFR; n = 4 for PTPRJ and nuclei). **(E)** EV and WT cells were treated with 10 µM gefitinib, 3 µM PHA665752, or DMSO for 3 days, and 10 ng/mL EGF and 50 ng/mL HGF were added for 15 minutes before lysis (n = 3 per condition). Immunoblotting was performed for the indicated proteins. **(F)** HSC3 EV and WT cells were treated with three growth factors concurrently (10 ng/mL EGF, 50 ng/mL HGF, and 0.1 ng/mL NRG1), or vehicle, for 24 hours and 18 hours before lysing. Immunoblotting was performed for the indicated proteins (n = 3 per condition). **(G)** HSC3 WT cells were treated with 10 ng/mL EGF, 50 ng/mL HGF, or 37.5 ng/mL NRG1, or vehicle, 24 hours and 18 hours before lysing (n = 3 per condition), and immunoblotting was performed for the indicated proteins. Unless otherwise stated, one-way ANOVA and post-hoc Sidak’s multiple comparisons; not all comparisons shown; * p < 0.05, ** p < 0.01, *** p < 0.001, **** p < 0.0001, n.s., nonsignificant.

Given the strong ability for HER3 to promote PI3K activity (45, 46), we checked for an effect of HER3 depletion on AKT phosphorylation, anticipating that such an effect could be responsible for the proliferation changes. In EV and WT cells treated with HGF, HER3 loss suppressed AKT phosphorylation, but the effect was not apparent in G983L cells, presumably due to the elevated activity of the mutant PTPRJ (Fig. 8B). Unexpectedly, HER3 depletion also reduced PTPRJ expression in cells with ectopic expression, although the effect was only significant for G983L cells. The unexpected loss of PTPRJ expression with HER3 knockdown strengthened our conclusion of the importance of HER3 for regulating phenotypes. That is, the fact that a loss of PTPRJ with HER3 knockdown did not restore proliferation suggests that HER3 was a critical node regulated by PTPRJ. Given that PTPRJ was ectopically expressed under the control of an exogenous viral promoter, the regulation of PTPRJ expression by HER3 most likely occurred post-translationally.

To determine if the loss of AKT activity observed with HER3 knockdown could be responsible for the concomitant loss of PTPRJ expression, we inhibited PI3K using GDC-0941 in the WT cell background (Fig. 8C). PI3K inhibition significantly reduced AKT phosphorylation and led to a small, but statistically significant, loss of PTPRJ expression. To test if the loss of AKT activity with HER3 knockdown could also be responsible for reduced proliferation observed with HER3 knockdown, we again used the PI3K inhibitor (Fig. 8D). PI3K inhibition reduced proliferation only for G983L cells treated with HGF. Given the general discrepancies between conditions where HER3 knockdown or PI3K inhibition antagonized proliferation, we concluded that regulation of AKT activity was unlikely to be the only means by which PTPRJ exerted effects on cell phenotype via its regulation of HER3.

In addition to the ErbB family, HER3 is a substrate of other receptors, including MET (47–50). Accordingly, we asked if EGFR or MET inhibition would reproduce the effect of HER3 loss and limit PTPRJ expression. Treatment of WT cells with an EGFR or MET inhibitor indeed modestly but significantly suppressed PTPRJ expression, but only the EGFR inhibitor antagonized AKT phosphorylation (assessed in the EV background due to the higher baseline AKT phosphorylation) (Fig. 8E). Issues of differential treatment timings and cell backgrounds notwithstanding, one interpretation of these results is that PTPRJ expression is regulated downstream of RTKs via pathways other than AKT.

Given that inhibition of EGFR or PI3K or knockdown of HER3 reduced PTPRJ expression, we hypothesized that RTK activation would reciprocally stabilize PTPRJ expression. Simultaneous treatment of WT cells with EGF, HGF, and NRG1 (a ligand for HER3) elevated PTPRJ expression, but there was no measurable effect in EV cells, presumably because of the low background expression in that cell line (Fig. 8F). Reduced expression of EGFR in response to the growth factor cocktail provided evidence of EGFR activation and endolysosomal trafficking. To isolate the receptor most responsible for stabilizing PTPRJ expression, we treated WT cells with EGF, HGF, or NRG1 individually (Fig. 8G). Only EGF significantly increased PTPRJ expression. Contrary to the results observed in the presence of HGF, when HER3 was knocked down in the presence of EGF, PTPRJ expression was unaffected. Thus, EGFR activity stabilized PTPRJ expression in a manner independent of the HER3-regulated mechanism that we also identified. In aggregate, these results identify a ligand context-dependent role for HER3 in stabilizing PTPRJ expression that is overcome in the presence of EGF. More generally, these results identify an undocumented network of reciprocal regulatory interactions among PTPRJ and two members of the ErbB family of receptor tyrosine kinases.

### JNK regulates cell migration through CREB-independent pathways

PLSR models predicted that PTPRJ-regulated c-Jun N-terminal protein kinase (JNK) promoted migration and wound healing (Fig. 3C). To test this, we repeated scratch assays in the presence of HGF and IGF1 and with or without the JNK inhibitor SP60015 (Fig. 9A). JNK inhibition significantly suppressed wound closure for EV and WT cells but not for G983L cells, presumably because wound closure was already slow in G983L cells. Among HSC3 groups treated with JNK inhibitor, G983L cells exhibited a significant decrease in wound closure only when treated with HGF. We interpreted these data to indicate that JNK was a PTPRJ-regulated pathway in multiple growth factor contexts that strongly controls cell migration.

**Figure 9.**
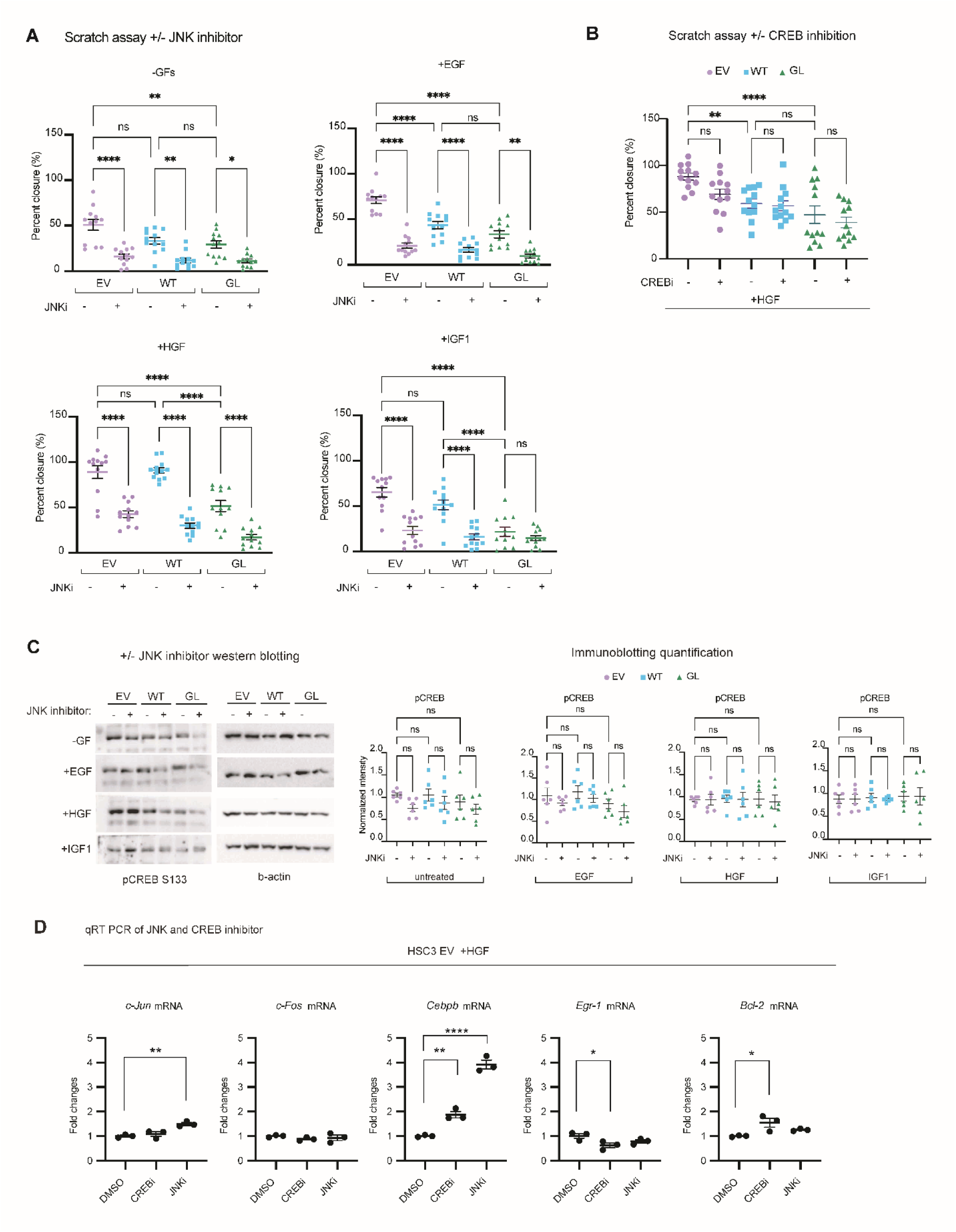
PTPRJ-regulated JNK signaling influences cell migration. **(A)** HSC3 cells were pre-treated with 20 µM SP60015 for 1 hour, then media was replaced with 0.1% FBS-containing DMEM with 10 ng/mL EGF, 50 ng/mL HGF, 10ng/mL IGF1, or no growth factors for the duration of a scratch assay (n = 12 per condition). EV = empty vector; WT = wild type; GL = G983L. **(B)** HSC3 cells were treated with 1 µM 666-15 or DMSO for one hour, and then media was replaced with 0.1% FBS-containing medium with 50 ng/mL HGF for the duration of the scratch assay (n = 12 per condition). One-way ANOVA with Sidak’s multiple comparisons test; * p < 0.05, ** p < 0.01, *** p < 0.001, **** p < 0.0001; n.s., nonsignificant. **(C)** HSC3 cells were treated with 20 µM SP60015 for one hour, and then media was replaced with 0.1% FBS-containing media with 10 ng/mL EGF, 50 ng/mL HGF, 50 ng/mL IGF1, or no additional growth factors, for 15 minutes before cells were lysed. Immunoblotting was performed for the indicated proteins. **(D)** HSC3 EV cells were treated with DMSO, 1 µM 666-15 or 1 µM SP60015 for 24 hours, and all were treated with 50 ng/mL HGF for 24 hours before mRNA extraction. qRT-PCR was performed for the indicated transcripts with two technical replicates (n = 3 per condition). Welch’s t-test; * p < 0.05, ** p < 0.01, *** p < 0.001, **** p < 0.0001, n.s., nonsignificant.

We next investigated the role of cyclic adenosine monophosphate (cAMP) response element binding protein (CREB) due to its proximity to JNK healing in the scratch PLSR model (Fig. 3) and because there is evidence of JNK-CREB crosstalk. JNK can phosphorylate CREB or proteins translated downstream of CREB’s role as a transcription factor, such as Bcl-2 (51, 52). In some settings, JNK inhibition surprisingly increases CREB phosphorylation and abundance of downstream transcripts such as *c-Fos* (53). JNK inhibition occurs through cAMP-mediated CREB regulation of downstream c-FLIP and MKP-1 (54) in some settings. Finally, the c-Jun target of JNK and c-Fos downstream of CREB dimerize (55).

Based on those inferences from the literature, we tested the ability of the JNK inhibitor to suppress CREB phosphorylation (Fig. 9B). None of the effects we measured across cell backgrounds or ligand treatments was significant. A purported CREB inhibitor failed to slow wound closure of any of the PTPRJ cell backgrounds (Fig. 9C). Note that scratch assays were performed in the presence of HGF based on the apparent ability of that growth factor to promote wound healing and the model-predicted role of CREB in that setting. To check the efficacy of the CREB inhibitor we used, we tested for potential changes of reported transcriptional targets of CREB in EV cells treated with HGF (Fig. 9D). We expected that *Cebpb* would at least be suppressed by CREB inhibition, because increased CREB activity has been found to upregulate *Cebpb* in the context of inflammation and tissue repair (56). JNK inhibition resulted in an unanticipated increase in *c-Jun* and *Cebpb* transcripts. CREB inhibition resulted in an unanticipated increase in *Cebpb* transcripts, mirroring the effect of the JNK inhibitor. CREB inhibition also slightly reduced *Egr-1* transcript abundance. These inferences should be pursued by follow-up immunoblotting experiments.

## DISCUSSION

We characterized the ability of PTPRJ to regulate signaling and phenotypes driven by distinct RTKs. The extensive list of PTPRJ-regulated signaling nodes we identified included EGFR and MET, both previously reported as PTPRJ substrates, and other phosphoproteins that have not been reported as regulated by PTPRJ to our knowledge, including HER2, HER3, IGF1R, JNK, CREB, and NFκB. Phosphoproteins were typically modulated by PTPRJ to different degrees depending on the growth factor context and were typically more strongly regulated by the G983L mutant than WT PTPRJ. For conditions without exogenous growth factors, PTPRJ ectopic expression reduced HER2 and HER3 phosphorylation. The exogenous growth factors made the dephosphorylation of EGFR, MET, and IGF1R with ectopic PTPRJ expression more pronounced, but the suppression of pHER2 and pHER3 by PTPRJ groups remained under all growth factor contexts. Whether this reflects a higher baseline phosphorylation of HER2 and HER3, enabling a difference to be apparent without growth factors, or some structural feature that enables PTPRJ to potentially access these ErbB receptors without a driving growth factor is unclear. Phenotypic outcomes however were growth-factor dependent, with HGF being the most relevant growth factor promoting growth, adherence, and survival.

We dedicated substantial effort to probing the unexpected regulation and phenotypic importance of HER3 and an apparent HER3-PTPRJ feedback that was revealed through HER3 knockdown studies. There is relatively little information about specific HER3 phosphatases, but the type-2 RPTPs PTPRO and PTPRH and cytosolic PTPN9 are reported regulators (31, 57, 58). WT and substrate trapping PTPRO immunoprecipitate with HER3, leading to the conclusion that PTPRO complexes with HER3 independently of its catalytic domain (58). Interactions between PTPRH and HER3 were found through a screen of STM RPTP-ErbB interactions (31). This study also probed possible PTPRJ-HER3 binding within a PTP-RTK panel but found nonsignificant interactions for HER3 and PTPRJ in response to EGFR activation, suggesting that cell background and growth factor abundance may influence interactions (31). PTPN9 antagonizes HER3 phosphorylation without affecting EGFR phosphorylation (57). HER3 immunoprecipitates with WT PTPN9 but not a substrate trapping PTPN9, which was interpreted to indicate that PTP9N may be a HER3 substrate. These examples illustrate the complex and varied nature through which an RTK regulation by phosphatases may proceed.

Our identification of strong regulation of HER3 phosphorylation by PTPRJ adds another important effector of PTPRJ, whether HER3 is a substrate or indirectly regulated target. Due to its low intrinsic catalytic activity, HER3 is typically viewed as a pseudokinase that must partner with a more active RTK such as EGFR to form a signaling-competent dimer and regulate downstream effectors (44, 59). HER3 is also a substrate of amplified MET, and this interaction drives resistance of lung cancers to EGFR inhibitors (47, 59, 60). Given those points and recalling that HER3 has not been reported in substrate-trapping studies, it seems likely PTPRJ indirectly suppressed pHER3 in our studies through its regulation of EGFR or MET. The ability of PTPRJ to regulate HER3 phosphorylation strongly even in the absence of exogenous growth factors, however, argues against this model. This raises the possibility that another unidentified substrate modulates HER3 or HER3/RTK interactions.

We observed an unexpected reduction in the expression of ectopic PTPRJ with HER3 knockdown. Inhibiting the HER3 effector PI3K similarly suppressed PTPRJ expression. With 24-hour EGF treatment, HER3 knockdown did not produce the same reduction in the expression of ectopic PTPRJ, suggesting a potential ability for EGFR activity to stabilize PTPRJ expression. Treating cells with exogenous EGF alone confirmed this inference. Thus, the experiments we performed, motivated by model inferences, led us to reveal a PTPRJ-ErbB-MET-AKT feedback loop. Other potentially related RTK-PTP relationships have been reported. Lung cancer cells treated with gefitinib (an EGFR kinase inhibitor) displayed reduced expression of PTP-PEST, a known EGFR phosphatase (61, 62). It is important to remember that the transgenes encoding WT and G983L PTPRJ were under the control of a non-endogenous, viral promoter in the cells we engineered. Thus, it is likely the changes we observed in PTPRJ expression as a function of PTPRJ perturbation arose due to post-translational modifications rather than transcriptional regulation. Where such regulation might arise is unclear though. Unlike PTP-PEST (63), PTPRJ is not known to have a high-strength PEST domain that can stabilize its expression in response to phosphorylation.

The results of our study confirm prior inferences from our work that targeting the TMD of PTPRJ can be a useful way to modulate PTPRJ-dependent signaling in cancer cells. In contrast to pro-oncogenic phosphatase SHP2, which can be inhibited allosterically by SHP-099 (64) and various related compounds in clinical trials, we found that promoting the activity of tumor suppressor PTPRJ suppressed growth, migration, adherence, and survival under cytotoxic conditions. Other PTP-targeting drugs are being developed. For example, combined PTP1B-PTPN2 active-site inhibitor ABBV-CLS-484 is being investigated in phase I trials for NSCLC and HNSCC (#NCT04777994). Several PTP1B inhibitors are also being investigated for type 2 diabetes, a context in which there is frequently insulin resistance due to insulin receptor and IRS dephosphorylation (through which PTP1B is implicated), downstream insulin pathway aberrations, and many other mechanisms (65).

Our results suggest that therapeutic activation of PTPRJ could be particularly effective in the context of carcinomas driven by HER3, MET, and EGFR. HER3 and high HGF-dependent signaling were predicted to drive many of the phenotypes we measured, but PTPRJ G983L was positioned as the cell background best controlling growth, proliferation, and adherence. Uncontrolled growth, lowered adherence, and resistance to cytotoxicity are hallmarks of cancer. Our analysis pointed to PTPRJ G983L as a strong modulator of proliferation and adherence (i.e., measure of cell index) and thus could be relevant to contexts of higher EMT or metastatic carcinomas.

Computational modeling led us to test hypotheses that would otherwise have been unintuitive. For example, because the phosphorylation of all ErbB receptors was modified by PTPRJ expression differences, it would have been unlikely to pursue HER3 as a specific regulator of PTPRJ-controlled signaling and phenotypes simply by inspecting the Luminex data. The predicted JNK-CREB axis would also have been unlikely to be tested without the PLSR model. Of course, PLSR and related modeling approaches have limitations. A classical limitation of these non-mechanistic modeling approaches is the possibility of pointing to a correlation without causation. Both of our major modeling validation efforts pointed to this. Even though JNK was validated as highly relevant to wound healing, the validity of the predicted role of CREB and a possible JNK-CREB axis remains unclear. A larger network of proteins may be involved in a possible JNK-CREB axis, and regulation via CREB may occur in a JNK-independent manner. Even with these ambiguities, the hypothesis generation that arises from inspection of a PLSR or other related model type frequently leads to important but unanticipated mechanistic understanding.

## ACKNOWLEDGEMENTS

This work was supported by NIH/NIGMS R01-GM139998, the UVA Biotechnology Training Program T32-GM136615, and NCI P30-CA044579.

---

**Figure Supplement 1.**
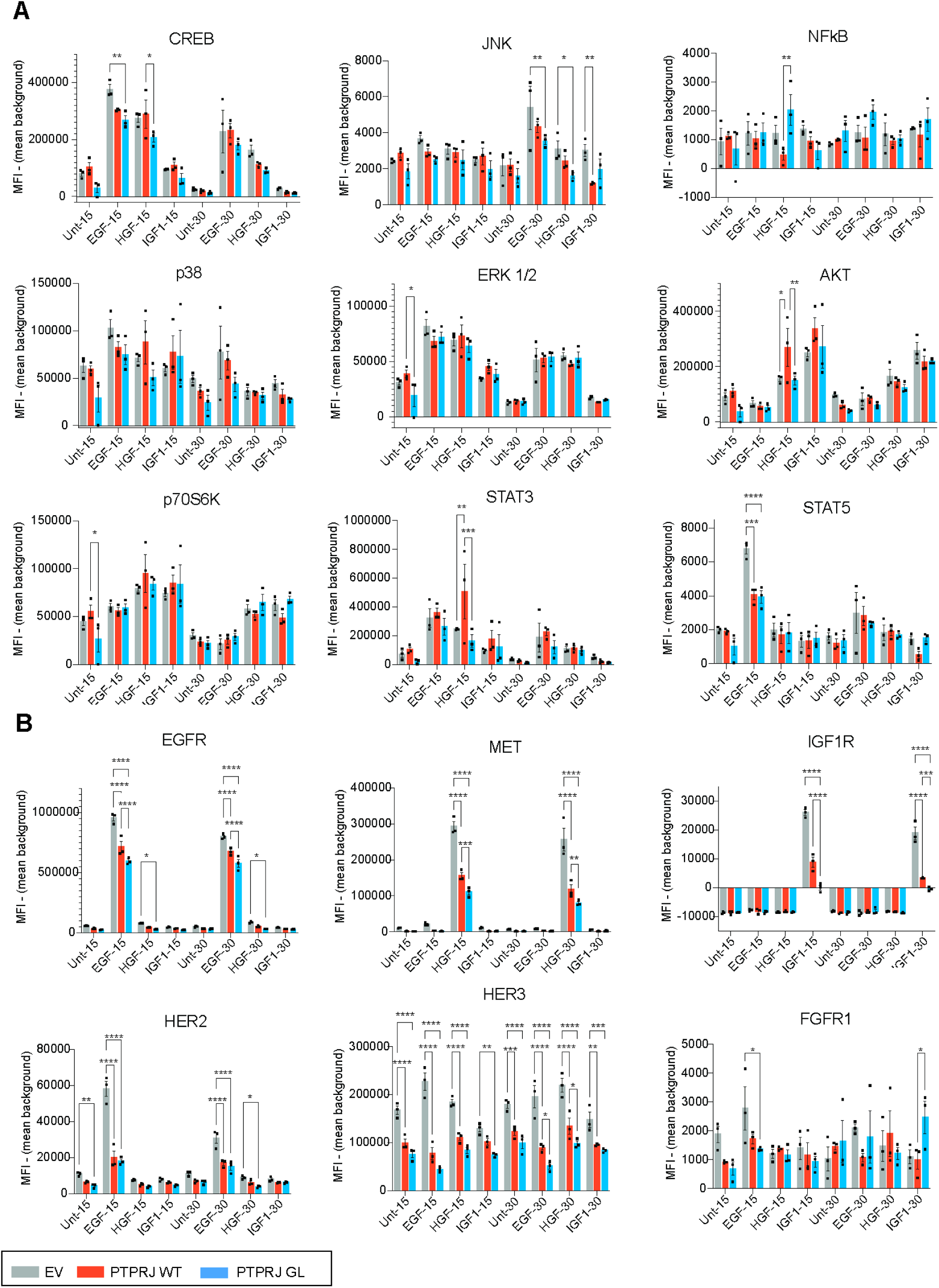
Summary of Luminex data. **(A)** Multiplexed phosphoprotein measurements from the Milliplex Multipathway kit (cat# 48-680MAG). **(B)** Multiplexed measurements receptor tyrosine kinase proteins from the Milliplex RTK kit (cat# 48-671MAG). Two-way ANOVA with Tukey’s multiple comparisons test; * p < 0.05, ** p < 0.01, *** p < 0.001, **** p < 0.0001, n.s., nonsignificant.

**Supplemental Table 1.**
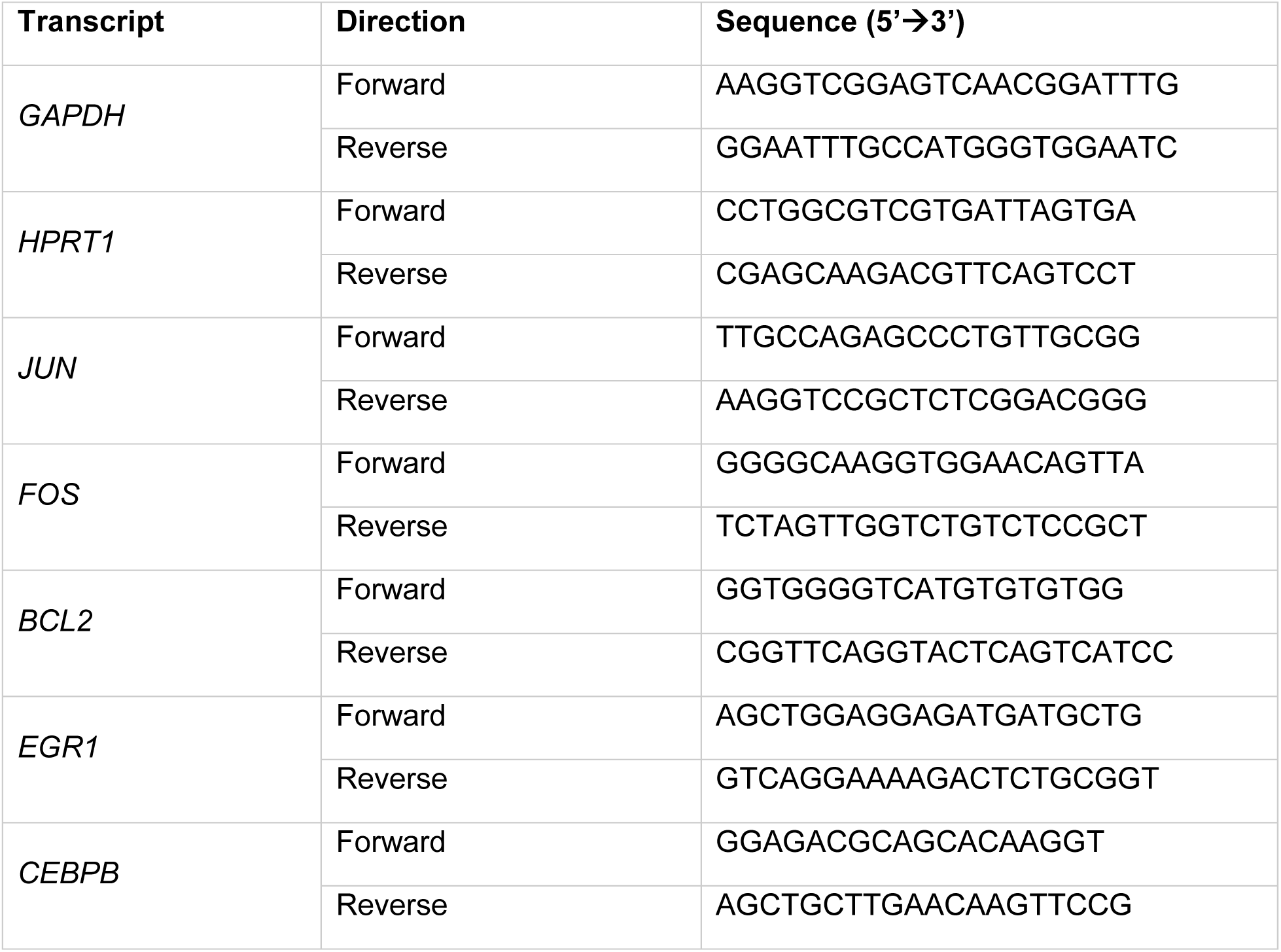
qRT-PCR primers.

**Supplemental Table 2.** Immunofluorescence antibodies.

| **Antibody** | **Catalog number** | **Dilution** |
| --- | --- | --- |
| PTPRJ mouse | Santa Cruz Biotechnology #sc-376794 | 1:100 |
| HER3 rabbit | Cell Signaling Technology #12708 | 1:100 |
| pAKT S473 rabbit | Cell Signaling Technology #9271 | 1:200 |
| EGFR rabbit | Cell Signaling Technology #4267 | 1:100 |
| Goat Anti-rabbit 647 | Invitrogen A21245 | 1:750 |
| Donkey Anti-mouse 546 | Invitrogen A10036 | 1:750 |
| Goat anti-mouse Alexa Fluor 647 | Invitrogen #A32728 | 1:750 |
| Goat anti-rabbit Alexa Fluor 546 | Invitrogen #A11035 | 1:750 |

